# PriMAT: A robust multi-animal tracking model for primates in the wild

**DOI:** 10.1101/2024.08.21.607881

**Authors:** Richard Vogg, Matthias Nuske, Marissa A. Weis, Timo Lüddecke, Elif Karakoç, Zurna Ahmed, Sofia M. Pereira, Suchinda Malaivijitnond, Suthirote Meesawat, Derek Murphy, Julia Fischer, Florentin Wörgötter, Peter M. Kappeler, Alexander Gail, Julia Ostner, Oliver Schülke, Claudia Fichtel, Alexander S. Ecker

## Abstract

1. Detection and tracking of animals is an important first step for automated behavioral studies using videos. Animal tracking is currently done mostly using deep learning frameworks based on keypoints, which show remarkable results in lab settings with fixed cameras, backgrounds, and lighting. However, multi-animal tracking in the wild presents several challenges such as high variability in background and lighting conditions, complex motion, and occlusion.
2. We propose a multi-animal tracking model, PriMAT, for nonhuman primates in the wild. The model learns to detect and track primates and other objects of interest from labeled videos or single images using bounding boxes instead of keypoints. Using bounding boxes significantly facilitates data annotation and robustness. Our one-stage model is conceptually simple but highly flexible, and we add a classification branch that allows us to train individual identification.
3. To evaluate the performance of our model, we applied it in two case studies with Assamese macaques (*Macaca assamensis*) and redfronted lemurs (*Eulemur rufifrons*) in the wild. We show that with only a few hundred frames labeled with bounding boxes, we can achieve robust tracking results. Combining these results with the classification branch for the lemur videos, our model shows an accuracy of 84% in predicting lemur identities.
4. Our approach presents a promising solution for accurately tracking and identifying animals in the wild, offering researchers a tool to study animal behavior in their natural habitats. Our code, models, training images, and evaluation video sequences are publicly available^1^, facilitating their use for animal behavior analyses and future research in this field.

## 1 Introduction

Automated analysis of actions and social interactions of free-ranging animals is essential for advancing behavioral research, enabling the study of complex behaviors in natural settings. Video recordings are a valuable tool for studying animal behavior, but traditional manual annotation of videos is time-consuming and labor-intensive. In the context of primate behavior research, large-scale annotated datasets have recently enabled the development and training of deep learning based models (Bala et al., 2020; Brookes and Burghardt, 2020; Labuguen et al., 2021; Yu et al., 2021; Desai et al., 2023; Yang et al., 2022; Ng et al., 2022; Ma et al., 2023). There are models which show promising results on several computer vision tasks in the wild, such as individual identification (Schofield et al., 2019; Bain et al., 2019; Schofield et al., 2023; Paulet et al., 2024), pose estimation (Wiltshire et al., 2023; Marks et al., 2022), object detection (Roy et al., 2023) and tracking (Koger et al., 2023; Pineda et al., 2023). However, the vast majority of frameworks used to analyze animal behavior are designed to work in lab settings and rely on pose estimation as an intermediate representation (Perez and Toler-Franklin, 2023; Vogg et al., 2024). Three of the most used frameworks for such markerless pose estimation are DeepPoseKit (Graving et al., 2019), DeepLabCut (Mathis et al., 2018; Lauer et al., 2022), and SLEAP (Pereira et al., 2022). They allow for multi-animal pose tracking and have user-friendly interfaces that help with annotation and inference. The introduction of these frameworks has contributed to great progress in the area of behavioral analysis.

As valuable as these tools are in the lab, their utility for applications of tracking animals in the wild is limited (Mathis et al., 2021; Perez and Toler-Franklin, 2023). Recording animals in their natural habitat presents challenges such as occlusion, diverse backgrounds, and varying lighting conditions. Similar appearances between individuals and sudden rapid movements add complexity to the task. We propose a different approach as a step towards analzying individual actions and social interactions in videos of wild primates. Instead of keypoints as an intermediate representation of the objects, our model outputs bounding boxes for all animals and relevant objects in the scene (Fig. 1).

**Figure 1:**
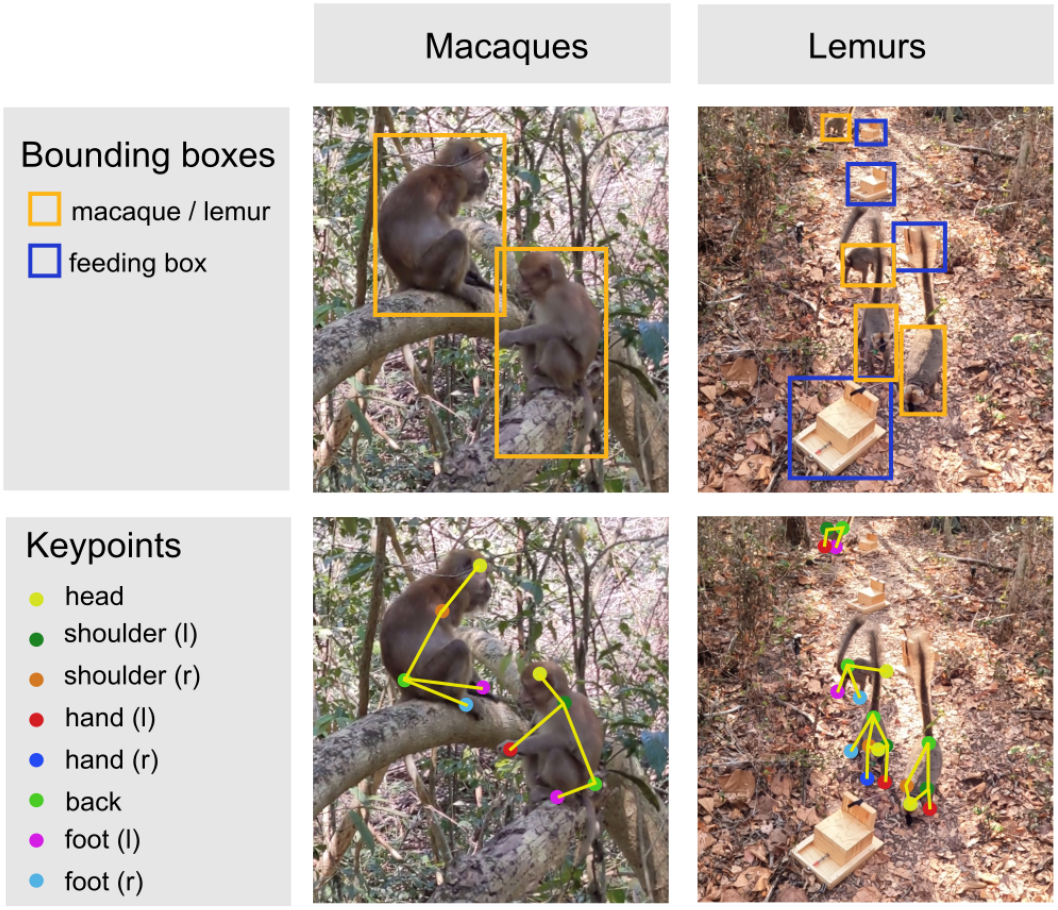
Bounding box based detection vs. keypoint detection on different datasets. Bounding box based tracking has advantages over keypoint tracking when used as a starting point for automated behavior analysis, such as reduced labeling time, higher tracking robustness for videos from the wild, and universal extensibility to other objects of interest.

We see three main reasons why object detection via bounding boxes should be a starting point to consider for many applications. First, currently, there are few successful applications of the above-mentioned keypoint-based frameworks on videos from the wild. DeepWild showed that DeepLabCut can learn from around 2000 labeled frames from videos of animals in the wild; however, the results on unseen videos leave room for improvement (Wiltshire et al., 2023) and more labels would be necessary to enhance generalization. A study conducted on horse images showed the limited capability of keypoint-based models to generalize to new, unseen horses (Mathis et al., 2021). A recent survey of keypoint-based methods for animal action classification found that most models do not generalize well to in-the-wild conditions (Perez and Toler-Franklin, 2023). For humans, pose tracking with variable backgrounds works (Xiao et al., 2018; Xu et al., 2022). The difference here is that diverse, large-scale datasets exist and substantial human labor has been put into annotating them (Ionescu et al., 2013; Andriluka et al., 2018). For most animal species, neither large-scale datasets with broad enough coverage exist, nor has an equivalent amount of labor been invested into annotations.

Second, using an object-detection framework built on bounding boxes may be preferable because keypoints require significantly more time to annotate than bounding boxes. Wiltshire et al. (Wiltshire et al., 2023) reported that they needed around two hours to label 18 keypoints for all animals in 25 frames. Their videos were chosen to have not more than seven individuals present. We annotated around 200 frames with bounding boxes in the same time.

Third, many questions in animal behavior research, especially in the wild, simply may not require keypoints but instead focus on actions and interactions which can directly be extracted from the image information inside the bounding box.

In this paper, we propose PriMAT, a conceptually simple **Pri**mate **M**ulti-**A**nimal **T**racking model. It builds on the FairMOT model (Zhang et al., 2021) which is trained on individual images and thus does not require entire video sequences to be labeled. Compared to keypoint labeling, it reduces annotation effort further, as bounding boxes can be drawn relatively quickly on a few hundred frames, which is enough for transfer learning to other primate species or camera angles. We added a branch to our model that can learn classification tasks, such as individual identification, on top of the tracking results. We present two use-cases for our model, by applying it to videos of Assamese macaques and redfronted lemurs in the wild. The models learned to detect macaques and lemurs from different camera angles and in videos with highly variable backgrounds and lighting conditions. Additionally, we demonstrate two extensions to the base model: First, multi-class tracking allows to detect other objects of interest in the video, which is a crucial step in analyzing interactions. We used it to detect lemurs and food boxes that had been placed on the ground as part of a behavioral experiment.Secondly, we used the classification branch to identify individual lemurs.

## 2 Methods

### 2.1 Data

Transfer learning is an efficient way to reduce annotation effort by starting with a base model which has been pretrained on a larger dataset and fine-tuning with a few hundred annotated frames from the domain of application. We experimented with four pretraining datasets (see below) and annotated 500 frames from videos of each our target species to showcase the applicability of the base model to macaques and lemurs (Fig. 2A). We used tight modal bounding boxes, i.e. boxes encompassing the extreme visible pixels of each individual.

**Figure 2:**
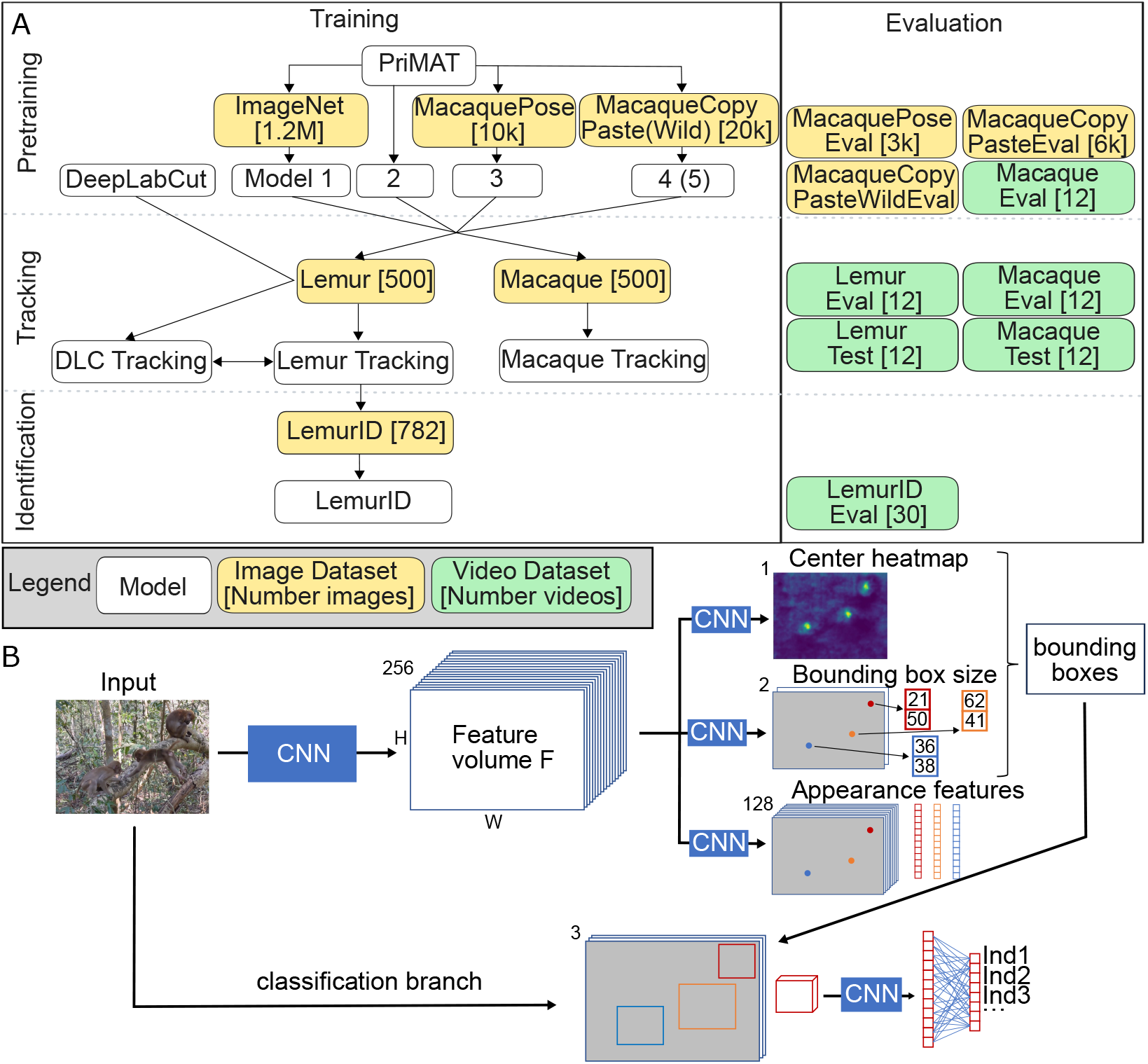
(A) Structure of our datasets and experiments. All models were trained with image datasets, while the performance was evaluated primarily on videos. We compared different datasets for pretraining, and applied them to two use cases: videos of redfronted lemurs and Assamese macaques. We compared our bounding box based approach with DeepLabCut (Lauer et al., 2022). Finally, we demonstrated the use of our additional classification branch for individual identification. (B) Overview of the model architecture: Input images are processed by a Convolutional Neural Network (CNN, in our case HRNet32, Sun et al. (2019)) into a feature volume. Different heads learn tracking-related tasks using two-layer CNNs. We omitted the offset head for simplicity. Afterwards, individuals are cut out and processed by the classification branch for individual identification. We used ResNet18 (He et al., 2016) as the CNN for individual recognition.

#### Pretraining

We compared ImageNet (Deng et al., 2009), MacaquePose (Labuguen et al., 2021) and two augmented versions of MacaquePose for pretraining. ImageNet is a standard dataset for pretraining, containing over 1.2M images from a variety of object classes. MacaquePose consists of 13083 images of macaques mostly taken from zoos. The dataset is completely annotated with instance segmentation masks and keypoints. We constructed bounding boxes from the extreme points of each segmentation mask. To improve generalization, we used the labeled instance segmentation masks from the MacaquePose dataset, copied the individuals and pasted them to backgrounds from ImageNet (Fig. 5A). The resulting images formed a dataset with high background diversity, which we called MacaqueCopyPaste. Additionally, we created MacaqueCopyPasteWild, a dataset where we pasted the monkeys on background images taken in the Phu Khieo Wildlife Sanctuary in Thailand. We used 80% of each pretraining dataset for training the model and kept the remaining images to evaluate the detection performance.

#### Assamese Macaques

Data on Assamese macaques were collected as part of a long-term project on the behavioral ecology of wild Assamese macaques in the Phu Khieo Wildlife Sanctuary, North East Thailand (Schülke et al., 2011). Data were collected on approximately 250 individually recognizable subjects of different age-sex classes organized in up to five large multi-male/multi-female social groups ranging in size from 40 to 73 individuals (unpublished data). The habitat consists of dense hill evergreen forest, and the monkeys spend the vast majority of their time in the trees, reducing visibility and creating difficult video-recording conditions. The videos were hand-recorded with resolutions of 1920 *×* 1080 using a GoPro Hero 10 camera and 3840 *×* 2160 using a Google Pixel 4 camera. More than 300 video snippets have been obtained so far. The videos consist of sequences that are challenging for a tracking model, as they can contain motion blur through camera or animal movement, severe occlusion, and high variability in backgrounds and lighting conditions.

We manually annotated the bounding boxes on 500 frames randomly sampled from the macaque videos. To extract meaningful samples, we used the model pretrained on MacaquePose, which yielded an approximate count of individuals in each video. Then, we sampled images from the macaque videos according to the macaque count, i.e. we chose more examples from crowded scenes and less from scenes with lone individuals. We used the software VoTT^2^, a user-friendly tool that can be used locally for annotating images to label single images for training.

We annotated 12 sequences, each one 12 seconds long, frame-by-frame with 30 frames per second, to evaluate the tracking performance with different hyperparameters and settings. Similarly, we annotated 12 sequences of the same length to form a test set, which can be used for benchmarking against our method. Validation and test videos were taken from experiments that were not used to extract training images. The sequences had varying levels of environmental complexity, and each included some motion or occlusion. We used the CVAT labeling software^3^ to label the evaluation videos. For video annotation, CVAT offers the advantage of automatically interpolating bounding boxes between frames, reducing the need for manual annotation when consecutive frames remain unchanged. This saves time compared to VoTT, which requires more manual input for frame-by-frame labeling. Each monkey in every frame was annotated if the annotator was able to recognize it as a monkey in the still image. If a monkey was temporarily occluded, it was labeled with the same ID after reappearing.

#### Redfronted Lemurs

We recorded videos during a social learning experiment in Kirindy Forest, Madagascar, that involved four groups of redfronted lemurs, with group sizes ranging from five to eight individuals. In each lemur group, we presented four feeding boxes in each experiment and repeated experiments ten times over the course of three months. The duration of each experiment varied from 10 to 30 minutes. We placed eight GoPro Hero 10 cameras to record the interactions of individuals with the feeding boxes from different angles (Fig. 3A). All cameras recorded videos with a resolution of 1920 *×* 1080 and 30 frames per second. Four cameras were placed to record the opening mechanism of each box from a distance of one meter (Fig. 3B). Two cameras mounted on nearby trees were used to capture a top-down view of pairs of boxes (Fig. 3C). Additionally, we placed two cameras on tripods to capture an overview of the experimental set-up from two opposite sides (Fig. 3D). Each video was treated independently by the model, without taking into account input by the other perspectives. We did, however, train one model to work well on all camera perspectives, instead of training three specialized models.

**Figure 3:**
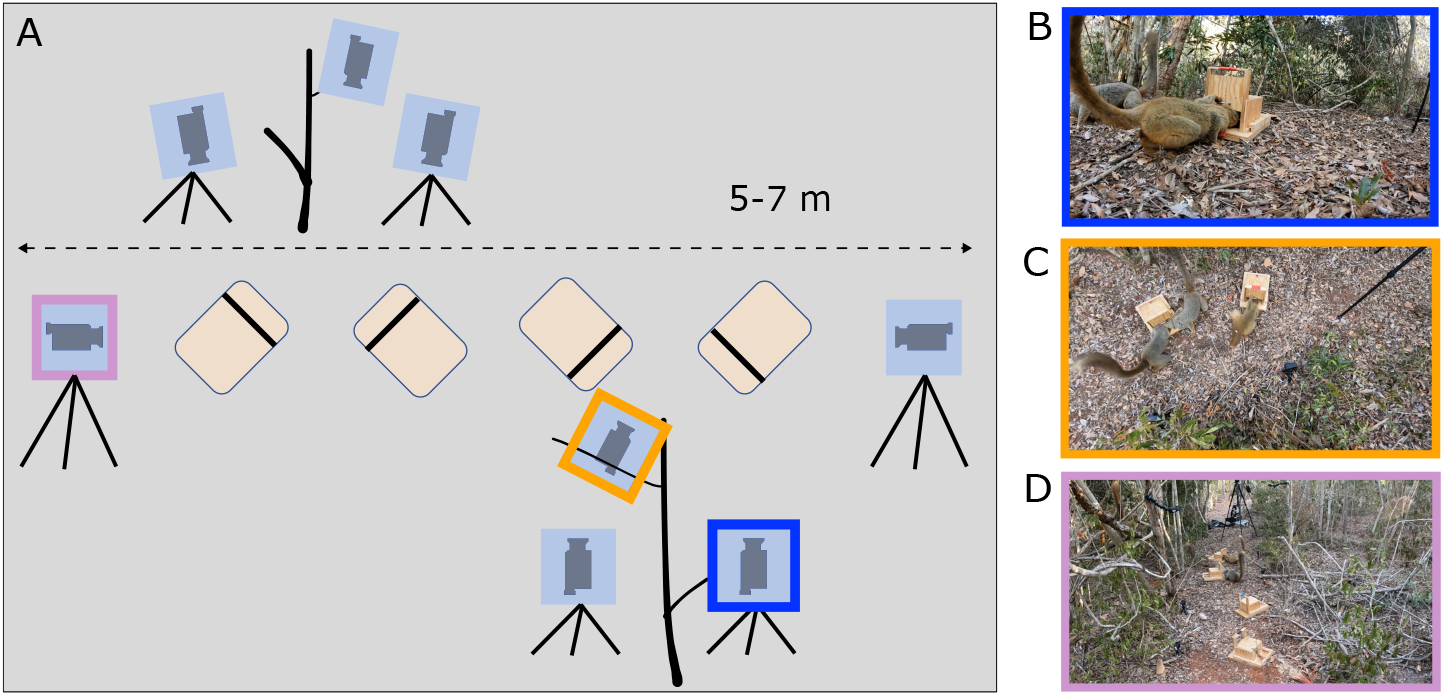
(A) Setup for social learning experiments: Four feeding boxes were placed on the ground and eight cameras were filming from three different perspectives ((B) close, (C) top, (D) far).

We recorded eight videos from different perspectives for each of the 40 experimental sessions. Additionally, we conducted pre-experiment sessions with open boxes so the lemurs could get used to the experimental setup. We used a subset of 50 videos from the pre-experiment sessions to obtain our training set. As the close-up view showed the most variability in lemur appearance, we used 24 videos from this view and 13 videos each from the far view and top views. In each video, we determined the longest segment that continuously contained at least one lemur, and randomly extracted 10 frames from each of these segments. The 500 resulting frames formed the training set. In these images, we annotated all lemurs by manually drawing the bounding boxes using VoTT^4^. We decided to exclude the lemur tails from the bounding boxes, as they would otherwise occupy a relatively large proportion of the enclosed space and lead to fast changes in bounding box size and location from frame to frame. Additionally, we labeled all feeding boxes in the videos to showcase the multi-class capabilities of our model.

For validation, we labeled bounding boxes for all lemurs and feeding boxes in 12 video sequences of length 12 seconds completely frame-by-frame (30 frames per second) to evaluate the tracking performance of the model. We chose four videos from each camera perspective of the experimental sessions, to avoid any overlap with the training images. We annotated a further set of 12 videos, which can be used for benchmarking against our method.

#### Individual Identification

All lemurs wore collars with tags of different shapes and colors to identify individuals. To train the model on the individual identification/classification task, we manually selected 350 frames from the close-up videos of one of the groups with six individuals as follows. Using 40 videos from this group, we extracted frames in which the collar of at least one individual was clearly visible. At maximum, we selected 25 frames from a single video to ensure diversity in background and lighting conditions. Again, we used VoTT to annotate the bounding boxes, this time labeled with each lemur’s identity. If any lemurs were present in these frames whose collars were not visible, we annotated them with the label “Unsure”.

To evaluate the identification performance on temporal tracks, we selected ten close-up camera videos and extracted three one-minute video sequences from each. We manually chose sequences in which the lemur tracking model had no ID switches in order to have an unambiguous ground truth with one individual ID per track.

#### Ethical approval

This study adhered to the Guidelines for the Treatment of Animals in Behavioral Research and Teaching (Buchanan et al., 2012) and the legal requirements of the countries (Madagascar, Thailand) in which the work was carried out. The Malagasy Ministry of the Environment, the Mention Zoologie et Biodiversité, Animale de l’Université d’Antananarivo and the CNFEREF Morondava approved and authorized this research (036/24/MEDD/SG/DGGE/DAPRNE/SCBE.Re). Data collection in Thailand was authorized by the Department of National Parks, Wildlife and Plant Conservation (DNP) and the National Research Council of Thailand (NRCT) (permit numbers 0401/11121 and 0401/13688).

### 2.2 Multi-Animal Tracking of Primates

PriMAT is based on the FairMOT (Zhang et al., 2021) architecture, a popular multi-object tracking model initially developed for tracking pedestrians. Our model is structured into three parts (Fig. 2B): *(1)* a convolutional backbone used for feature extraction, *(2)* multiple heads that perform different tasks for tracking, and *(3)* a classification branch. More detailed information about backbone, heads and association methodology can be found in the Supplementary Material.

Due to the existing datasets (Shao et al., 2018; Ionescu et al., 2013) and benchmarks (Milan et al., 2016; Dendorfer et al., 2020; Sun et al., 2022) for multi-object tracking, FairMOT, like most tracking models, was designed for tracking human pedestrians. However, there are large differences between humans and nonhuman primates, as well as among nonhuman primate species, in appearance, motion patterns, and acceleration. Therefore, we adapted the tracking strategy of our model to make it more suitable for the appearance and motion of monkeys and lemurs as outlined below.

Pedestrians in city scenes are generally easily distinguishable from the background Sun et al. (2022). This makes them easier to detect and track than nonhuman primates in their natural habitat, especially when the primates are in non-standard poses or highly occluded. Therefore, we lowered the confidence threshold for the center heatmap to values between 0.01 to 0.04 depending on the species, compared to the original 0.4 used for pedestrian tracking. This adjustment was necessary to maintain track continuity, especially in cases where the animals were moving. As low detection thresholds introduce potential false positive detections, we added a threshold to prevent new, unmatched boxes from starting a new track if they have too much overlap with already existing tracks. Additionally, we removed detections less than five pixels away from the image border, as most false positives were found in these areas. The appearance of pedestrians is highly variable between individuals, and therefore, the original FairMOT model uses only learned appearance features for when associating existing tracks with new detections. The unmatched tracks and detections are then compared using intersection-over-union (IoU). Nonhuman primates have much less inter-individual variation in appearance, thus, for our use-cases, location is a more important predictor than appearance. For this reason, we use a linear combination of appearance and location similarity in one step to calculate association.

Nonhuman primates also exhibit greater variability in acceleration and movement patterns compared to pedestrians. Tracking models typically use a Kalman filter to predict the location of objects from frame to frame. The Kalman filter assumes linear motion of objects and predicts the position of an object based on the velocity over the previous frames, which may not be appropriate for tracking nonhuman primates. For instance, when jumping, a lemur moves with high velocity, which then drops close to zero once the animal lands. This kind of movement can cause problems when calculating the association (Fig. 4). To avoid problems with fast motion changes, we added a conditional second stage to the association procedure. If there is no match at the location predicted by the Kalman filter, we additionally check for matches at the location of the last detection.

**Figure 4:**
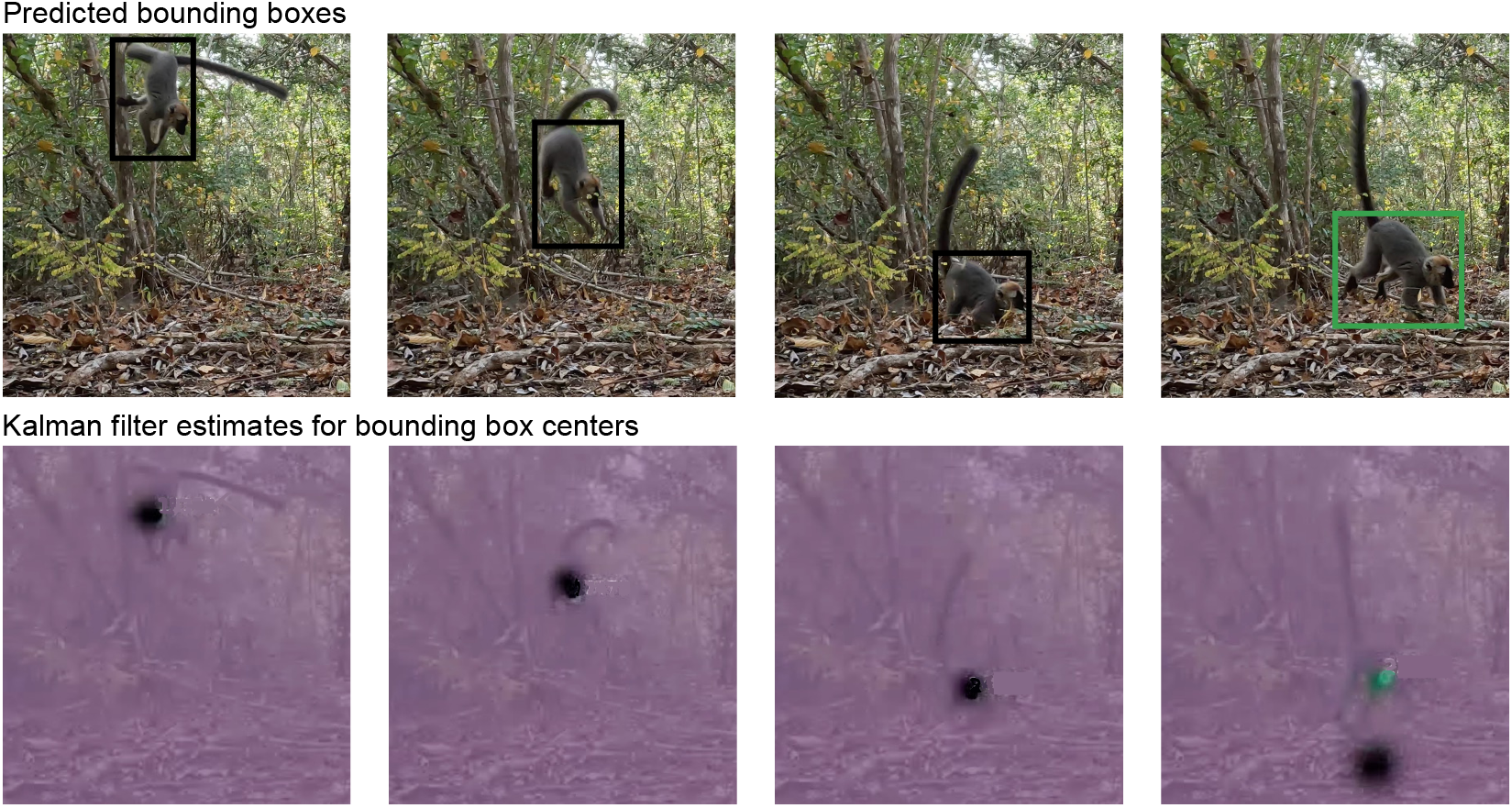
Problem during a jump: The Kalman filter predicts the location of the lemur track based on past velocity. The detection in the last frame cannot be matched to the lemur track. Top row: Detected bounding boxes and assigned tracks identified by the color of the bounding box. The track of the jumping lemur is lost in the last frame (black) and a new track is instantiated (green). Bottom row: Kalman filter predictions for the expected position of the lemur. The Kalman filter assumes linear motion and predicts a different position when the jump ends (black). The newly instantiated track starts with default parameters for the Kalman filter prediction (green).

### 2.3 Individual identification

To demonstrate how the tracking model can be extended, we added a new branch to the model and trained it to identify individual lemurs. It is important to note that we were not aiming for frame-wise correct identification, which would have been infeasible, as the lemurs were frequently in positions where identification was impossible. Instead, we used the tracks to arrive at a majority vote on which lemur is visible in each sequence of detections. This required the model to be confident in situations where the lemur is clearly identifiable and not be overconfident in situations where the decision could not be made. The final decision for which individual is predicted for a track was made by weighting and summing the confidences for each frame in a majority vote.

We trained the classification branch for individual identification using a ResNet18 (He et al., 2016) model pretrained on the ImageNet (Deng et al., 2009) dataset. We fine-tuned the network with the individual lemur images, which we cut out from the training images. As ResNet18 expects square images as input, we used squared bounding boxes to avoid squeezing the rectangular bounding boxes that we had previously annotated into squared shapes and introducing biases to the training set. We achieved the squared bounding boxes by increasing the size of the shorter sides of the previously annotated bounding boxes to match the longer sides and then cropped and resized them to 224 *×* 224 pixels. To account for the shift from the perfectly annotated bounding boxes used during training to slightly imperfect bounding boxes predicted during inference, we applied random jitter of up to five pixels to the bounding boxes and random zoom increasing the width and height up to 20% during training. These values were chosen based on a randomized search over the hyperparameter space.

We trained the model with 315 hand-selected images of six individuals. The images were extracted from videos in situations where at least one individual was identifiable, either by their face or collar. After training, we used the model to acquire more training data, by applying it to four additional recordings of experiments and reviewing its predictions per track. We then created a second training set by randomly sampling 20 frames per track and selecting images manually, only including those that had no errors and were not near duplicates of other selected images. This process resulted in 782 new training images. Additionally, we used a small validation set of 35 hand-selected images for selecting the model via early stopping and evaluated the models on 30 one-minute-long snippets.

An alternative approach to using an independent model as described above is to add a new head directly to the backbone to learn identification jointly with the tracking task. We compared both model architectures and their performance in lemur identification.

### 2.4 Evaluation and metrics

#### Tracking

To evaluate how well our model learned to track individual primates, we applied it to the test set which consisted of 12 densely annotated video sequences (see Section 2.1 on datasets).

We evaluated the performance of our model with several metrics commonly used for multi-object tracking benchmarks. The most important metric is considered to be Higher Order Tracking Accuracy (HOTA) (Luiten et al., 2021), as it gives equal importance to detection and association. Additionally, we report Multi-Object Tracking Accuracy (MOTA) (Bernardin and Stiefelhagen, 2008) and IDF1 (Ristani et al., 2016), two traditional metrics for easier comparison to other tracking models.

#### Comparing the detection performance of bounding box and keypoints

We used a simple metric to compare detection performance from the bounding box and the keypoint tracking approaches, as we only had bounding box ground truth labeled for complete video sequences. For each ground truth bounding box, we determined whether there was a predicted bounding box with an intersection-over-union (IoU) value of at least 0.5, a standard threshold to decide whether a predicted bounding box should be considered to represent a ground-truth bounding box based on the amount of overlap between them (Lin et al., 2014). We then counted the number of predicted bounding boxes that overlapped sufficiently with each ground truth box (i.e. the number of correct detections). Similarly, we counted how often there were at least two ground-truth keypoints within a predicted bounding box. As the keypoints could be close to the box border, we also counted the keypoints that were within the 1.5× buffered bounding box. The value of 1.5*×* was chosen as it is a generous extension of the bounding box to allow even imprecisely detected keypoints to count towards a correct detection. For both approaches, we reported the proportion of correct detections out of the total number of ground truth bounding boxes as the recall value for each video.

Additionally, we calculated the precision of each model as the proportion of correct detections based on predicted keypoints or bounding boxes, respectively, out of the total number of predictions from each video. We reported the F1 score in both cases,

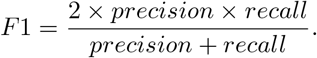

#### Detection

We evaluated the detection performance of bounding box based models using mean Average Precision (mAP), a classical metric for object detection tasks (Lin et al., 2014; Shao et al., 2018). In some cases we additionally reported AP50, a relaxed variant of mAP, where a bounding box counts as correctly predicted, if it has an IoU of at least 0.5 with a ground truth bounding box. Models which score high on AP50 but low on mAP are good in finding the presence of an animal, but have imprecise bounding boxes. For keypoint detection, we followed the metric suggested in DeepLabCut, and calculated the root-mean squared error for each detected keypoint to the closest ground truth detection (Lauer et al., 2022).

#### Individual identification

We performed the evaluation for individual identification on 30 one-minute-long videos containing 92 individuals. We evaluated track-wise results and used accuracy as a performance metric. Additionally, we generated confusion matrices of predicted vs ground-truth IDs to check for systematic errors between individuals.

### 2.5 Training details

PriMAT was implemented in PyTorch (Paszke et al., 2019) and trained with the Adam optimizer (Kingma and Ba, 2014). The training of the base model with MacaquePose (and the MacaqueCopyPaste datasets) was performed on four NVIDIA Quadro RTX 5000 GPUs for two days. We started with a learning rate of 5 *×* 10^*−*5^ and reduced it by a factor of 10 after 100 epochs, with a batch size of 8. For fine-tuning with 500 images for both lemurs and macaques, training on one GPU took six hours. The initial learning rate was 10^*−*5^. Early stopping could not be applied as we were training on images, but we were interested in tracking performance on videos.

For the individual identification task, we cropped the images to include only the bounding boxes surrounding lemurs in the dataset and stacked them afterwards, allowing training with a batch size of 128. The model was trained on a single NVIDIA Quadro RTX 5000 GPU, using early stopping with a patience of 10 epochs. Training time with a learning rate of 2 *×* 10^*−*5^ was one hour.

Additionally, we trained a model with DeepLabCut (Lauer et al., 2022) version 2.3.8 for which we annotated 33 keypoints per individual. We tested different hyperparameters for pcutoff (confidence for a detection) and IoU. Additionally, we compared DLCRNet ms5 (the data-driven individual assembly method which was recommended for improved performance in the multi-animal setting) and ResNet50 as backbones. We chose the best performing model with DLCRNet ms5, pcutoff 0.0 and IoU 0.6. All models were trained for 200,000 training iterations.

## 3 Results

With our multi-animal tracking model PriMAT, we were able to detect and track individual macaques and lemurs in videos from the wild, by only annotating bounding boxes on a few hundred frames. The model was specifically designed for good performance in primate videos, with solutions for non-standard poses, occlusion, similar appearance, and jumps (see methods). We evaluated the tracking performance of our model with annotated video data from Assamese macaques in Thailand and redfronted lemurs in Madagascar.

### Copy-paste data augmentation

We experimented with a simple copy-paste strategy for improved generalization to variation in the backgrounds. Previous work has shown that copying and pasting instances onto other backgrounds improved model performance in the area of instance segmentation (Ghiasi et al., 2021; Barreto et al., 2023) and object detection (Dwibedi et al., 2017; Yu et al., 2023) as it is an effort-efficient technique for creating realistic training images for any background. To train the base model, we used the images from the MacaquePose (Labuguen et al., 2021) dataset which show some background variability, however, they are mostly brown or grey (Fig. 5A) which is a domain shift when comparing to our videos in the green forest. To investigate this in more detail, we created MacaqueCopyPaste, a dataset with high background variability consisting of macaques pasted onto background images from ImageNet (Deng et al., 2009), and MacaqueCopyPasteWild with macaques pasted onto background images of the forest in Thailand.

**Figure 5:**
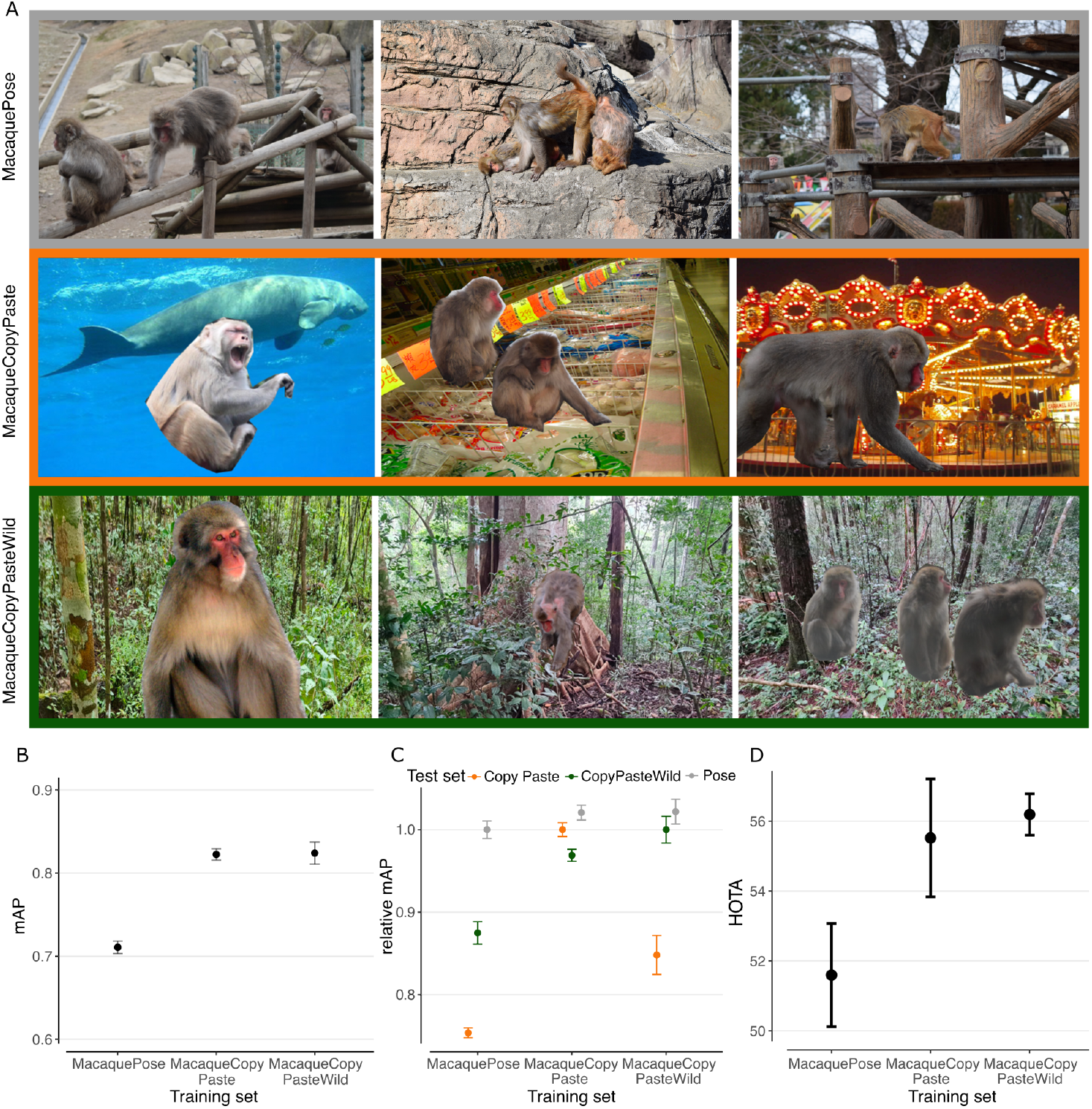
(A) Examples from MacaquePose (Labuguen et al., 2021) (top), MacaqueCopyPaste with ImageNet (Deng et al., 2009) backgrounds (middle), and MacaqueCopyPasteWild with backgrounds from Thailand (bottom). (B) Models trained on the copy-paste approaches yield higher results when tested *in-domain*, i.e. on their own test sets. This can probably be attributed to sharper edges and more saliency against the background. (C) Results of testing on different test sets relative to the in-domain performance from panel B. MacaqueCopyPaste detects macaques well on all three datasets. MacaquePose did not generalize well to other datasets. (D) Both copy-paste strategies outperform MacaquePose on the macaque validation videos. Error bars: Standard error of the mean from training three models with different random seeds.

We split the three datasets MacaquePose, MacaqueCopyPaste, and MacaqueCopyPasteWild into 80% training data and 20% test data and trained one model on each of them. We evaluated each model on the test set of all three datasets. When evaluating in-domain, we noticed that MacaqueCopyPaste and MacaqueCopyPasteWild showed better results (Fig. 5B), suggesting that these two datasets are “easier” than MacaquePose. Copying and pasting introduces sharper edges and often the animals are more salient against the background, compared to MacaquePose. We also evaluated how well the models performed across domains. While the models trained with MacaqueCopyPaste performed well on all three datasets, MacaquePose showed weaker results in the other domains. MacaqueCopyPasteWild performed well on MacaquePose but poorly on MacaqueCopyPaste (Fig. 5C). The tracking validation videos show similar backgrounds to the one MacaqueCopyPasteWild was trained on, which explains the better tracking performance. Both copy-paste approaches outperformed the models trained on MacaquePose (Fig. 5D).

We further investigated the ability of the base model pretrained on MacaqueCopyPaste to do transfer learning. We fine-tuned the base model with the 500 annotated images of lemurs and macaques. Pretraining with ImageNet (70.3 HOTA) was more beneficial than MacaqueCopyPaste (65.7 HOTA) for the lemurs (Table 1). The domain shift from the images of MacaquePose to the lemur videos is large, not only because the species look different, but the lemur experiments were filmed from different camera angles. For the macaques however, pretraining with MacaqueCopyPaste (66.1 HOTA) was slightly more beneficial than with ImageNet (64.6 HOTA). As we did a lot of hyperparameter tuning on the validation videos, we additionally provide 12 test sequences for each species. These can be used for benchmarking other methods against ours in the future. For the lemurs, the backgrounds and lighting conditions vary from experiment to experiment, however, the general setup is similar. The HOTA score on the test set (68.2) is similar to the validation set (70.3). For the macaques, the videos filmed from handheld cameras show higher variability. While the best performing model achieved 66.1 HOTA on the validation set, the score on the test set is 61.6 (see Table 1).

**Table 1:** Models were pretrained on different large-scale datasets and subsequently finetuned with 500 annotated images of lemurs or macaques. The tracking performance was evaluated on 12 validation video sequences. MacaqueCopyPaste pretraining resulted in improved results in macaque tracking. Pretraining on ImageNet showed better results for lemur tracking. Finally, we report test scores for the best validation setup (pretraining dataset, hyperparameters as detailed in the Supplementary material) for each species on 12 test video sequences.

|  | Lemurs |  |  | Macaques |  |  |
| --- | --- | --- | --- | --- | --- | --- |
|  | HOTA | MOTA | IDF1 | HOTA | MOTA | IDF1 |
| **Validation** |  |  |  |  |  |  |
| ImageNet (IN) | **70.3** | **81.5** | **88.1** | 64.6 | 75.0 | **83.7** |
| MacaqueCopyPaste (MCP) | 65.7 | 67.8 | 79.3 | **66.1** | **75.5** | **83.7** |
| no pretraining | 61.3 | 64.8 | 77.0 | 59.3 | 62.3 | 75.5 |
| **Test** |  |  |  |  |  |  |
| IN / MCP | 68.2 | 74.2 | 83.6 | 61.6 | 69.1 | 79.9 |

### Bounding box and keypoint based detection

In light of the studies mentioned in the introduction suggesting that models for keypoint estimation in the wild struggle with generalization, we performed a small experiment. To compare our model with keypoint-based tracking, we trained a DeepLabCut model (Lauer et al., 2022) to perform keypoint tracking on our lemur videos. We allocated five hours for annotation – the same time required to label 500 frames with bounding boxes - and annotated 33 keypoints per individual in 100 frames. These frames are a subset of the 500 frames used for training our model, ensuring comparable background variability by selecting fewer frames from all videos.

We evaluated the accuracy of keypoint tracking on seven selected keypoints (nose, head, neck, spine, start tail, mid tail, end tail) by annotating additional validation frames from the same 50 videos used for extracting training images. First, we observed that only 31% of the keypoints were detected as many ground-truth skeletons were completely missing or incomplete in the model predictions. Second, among the correctly detected keypoints, the median over the root-mean squared errors of all detections was 49.6 pixels. Of course, this value alone is difficult to interpret, as one pixel represents different distances in different views and we observed large variability between videos (Fig. 6B). However, in the lab, DLC models reach median test errors of under 5.25 pixels (Lauer et al., 2022) in all applications, which is almost a magnitude less than our model. The relatively large error value from our model was due to the prediction of multiple incomplete skeletons within an individual and skeletons that were split over two or more individuals. Next, we compared the performance of keypoint models and bounding box models in detecting the presence of a lemur (without localizing precise keypoints). As the data for each task involve a different level of granularity, and we only had bounding box ground truth labels for the validation videos, we designed a basic detection metric where two keypoints were needed to count as a successful detection. Our model outperformed the keypoint based method (Fig. 6C/D). In one video, which contained high-contrast shadows and therefore a greater need for generalization, the keypoint-based method was not able to detect any keypoints in the whole sequence. The performance of our bounding box method decreased on this challenging video compared to other videos, but it still resulted in a final F1 score of 0.62. The most notable difference between keypoints and bounding boxes was observed in the top view. For a qualitative comparison, we refer to the supplementary material.

**Figure 6:**
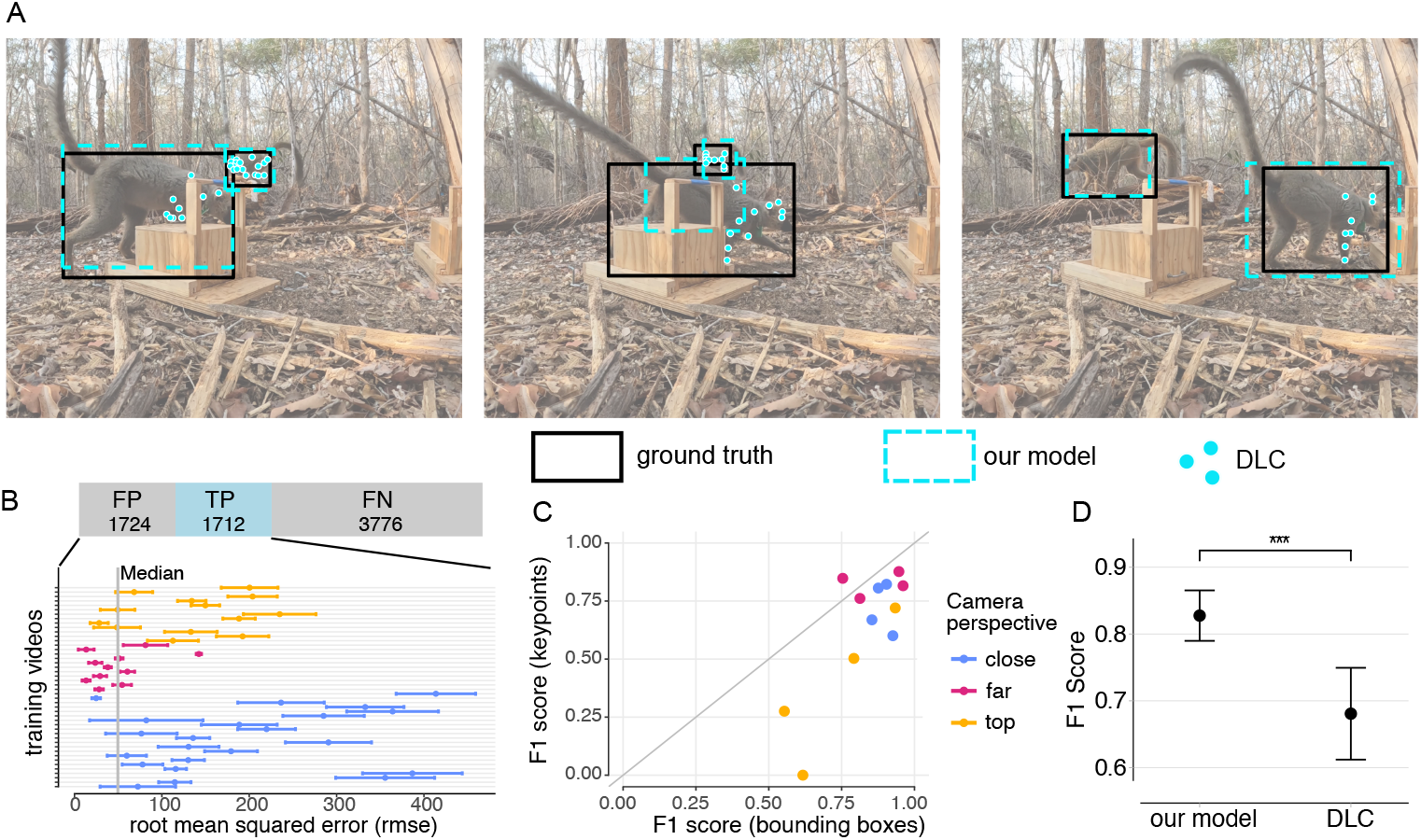
Comparison between our boxes based tracking model and a DeepLabCut (DLC) model (Lauer et al., 2022). Our model was trained with 500 images from three camera perspectives. The DLC model was trained using 33 keypoints, resulting in 100 frames. (A) A qualitative example where DLC does not detect most keypoints. (B) DLC missed many keypoints and shows large root-mean squared errors (rmse) for the predicted keypoints. (C, D) Performance difference on the 12 test videos. The evaluation metric was chosen in a favorable manner for DLC, as it counts a detection as positive when there are at least two keypoints correctly detected.

A model that outputs incomplete skeletons, many missing detections or incomplete tracks is of questionable utility for practical applications. To overcome these types of problem and have a robust and generalizing keypoint model, a lot more data is needed.

We performed an additional experiment in which we annotated only seven keypoints per individual and managed to annotate over 200 frames in the allocated five hours. However, this model performed worse than the one with 33 keypoints. This suggests that having a higher number of keypoints with cross-connections is more critical for performance than simply increasing the number of annotated frames.

### Individual identification

We added a classification branch described in Section 2.2 and applied it to identify individual lemurs in the videos. The final model had an identification accuracy of 84%. Additionally, we experimented with a different architecture. Instead of having two separate models for tracking and identification, we tested the possibility of adding a new head to the existing backbone. Table 2 shows the need to separate the two tasks, at least for the number of labels we used for training, as the model with shared backbone only reached an accuracy of 41%.

**Table 2:** Accuracy of models trained for individual identification. Having a shared backbone for tracking and individual identification showed poor results with the amount of data used (500 images for tracking, 782 images for individual identification). Feeding the detections into a second model for identification led to improved results.

| Architecture | Accuracy |
| --- | --- |
| Two separate models | 0.837 |
| Shared backbone | 0.413 |

### Transfer to other species

We trained a model jointly with the 500 images from the lemur and 500 images Assamese macaque videos to test its applicability to other settings or other primate species. We tested the model on videos of Barbary macaques (*Macaca sylvanus*) that were recorded as part of a decision-making experiment in Rocamadour, France (Bosshard et al., 2024), videos of Guinea baboons (*Papio papio*) ranging near the CRP Simenti Field Station in the Niokolo Koba National Park in Senegal (Fischer et al., 2017), and videos from the PanAf500 dataset, which includes chimpanzees (*Pan troglodytes*) and gorillas (*Gorilla spp*.) (Brookes et al., 2024). Qualitative results from applying the model to unseen videos can be found in the Supplementary material. We evaluated the detection performance with a 5-fold cross-validation. Applying the joint model without any further training showed good results for the Barbary macaques, even though the setting in these videos appears quite different from the ones in the videos from Thailand and Madagascar. The model was also relatively successful in detecting baboons, chimpanzees, and gorillas. However, particularly in the PanAf500 dataset, many individuals were not detected. This is likely because the species in this dataset (all great apes) have a very different appearance to lemurs and Assamese macaques, and the camera resolution is much lower than that of our videos. Performance increased significantly after fine-tuning with 100 frames from the target datasets. We observed high AP50 values which indicated that most individuals were correctly detected. The relatively lower mAP values indicated that the predicted bounding boxes did not fit perfectly around each individual (Table 3).

**Table 3:** Transfer results to other species. Evaluation metrics for models before and after fine-tuning with 100 frames from the target datasets

| Dataset | mAP | AP50 |
| --- | --- | --- |
| without training |  |  |
| Rocamadour | 0.55 | 0.83 |
| Guinea baboons | 0.22 | 0.56 |
| PanAf500 | 0.21 | 0.38 |
| finetuning (100 frames) |  |  |
| Rocamadour | 0.73 | 0.99 |
| Guinea baboons | 0.41 | 0.77 |
| PanAf500 | 0.47 | 0.81 |

## 4 Discussion

There are many potential paths towards automated behavior analysis using videos of animals in the wild. Most current methods in the lab use keypoint detection and tracking as a starting point, as those provide a valuable intermediate representation, namely the pose of the animal. However, so far, keypoint detection based models have struggled to generalize to highly diverse backgrounds in the wild. One solution could be to annotate vastly larger and diverse datasets; however, keypoint annotation is a very time-consuming task. Therefore, we proposed an alternative approach based on bounding boxes. We demonstrated that our deep learning based multi-animal tracking model, PriMAT, outperforms previous methods based on keypoints on videos from the wild while requiring significantly less labeling time. Our tracking model is based on the FairMOT architecture (Zhang et al., 2021), which was built to track pedestrians. However, pedestrians move almost exclusively in linear motion and wear distinctive clothing, which influenced the design choices for the model. We addressed primate-specific challenges in the design of our model, including non-standard poses, occlusion, and fast, non-linear motion.

### Limitations

We observed that PriMAT has problems with distinguishing two or more animals in situations where animals are sitting or moving very close together. A stronger detection model would be needed, potentially one that has been specifically trained with images that show animals sitting close together. One could also try to accomplish this with a copy-paste strategy and artificially create images with a lot of overlap between individuals. The model also had difficulties with detecting individuals when there was a lot of fast, non-linear movement and occlusion, such as in videos of juveniles playing. Our association strategy is rule-based and relies on a Kalman filter for linear movement (see Supplementary material). While adapting the association logic as done in our model can lead to further improvement, modern multi-object tracking architectures promise to overcome problems with challenging association by learning associations as part of their architecture instead of following hand-crafted rules (Zhou et al., 2022; Zhang et al., 2023). Our model processes each video frame independently without incorporating temporal information. Integrating temporal context could enhance performance by using motion patterns and improving consistency across frames. For the identification model, the resolution of the cut-out images of detected individuals strongly impacted the probability of a correct identification, as they are resized to 224 *×* 224 pixels regardless of their original size. We evaluated performance for different bounding box sizes and found that the model performed worse (45% accuracy) on tracks that were close to or smaller than 224 *×* 224 pixels originally compared to larger cut-out images (89% accuracy).

### Key findings and interpretations

We worked with pretrained base models that can be used as a starting point to learn to detect and track nonhuman primate species in different settings in the wild using only a few hundred labeled frames. Our lemur tracking model can track lemurs filmed from different camera perspectives and distances. As a step towards interaction classification, we also adapted the model to track other objects of interest, such as feeding boxes. Transfer experiments from macaques to lemurs suggested that ImageNet pretraining led to better results than pretraining on a large macaque dataset. When transferring to the macaque videos from Thailand however, datasets derived from MacaquePose yielded better results than ImageNet. We tested a strategy of copying and pasting macaques to different backgrounds, and our results showed that when doing transfer learning with a large domain shift (i.e. very different backgrounds in the training and test datasets), or a high background variability, the model trained with MacaqueCopyPaste (macaques on ImageNet backgrounds) was the best starting point. However, when there are similar backgrounds in the training and evaluation domains, such as with the models trained on our MacaqueCopyPasteWild dataset (macaques on backgrounds taken from the images from Thailand) and evaluated on the videos from Thailand, performance improved even further.

Our flexible model can learn additional classification tasks, such as individual identification, on top of tracking. With a total of 782 images of six individuals, we reached an accuracy of 84% in an individual identification task on unseen video tracks of lemurs wearing collars using a model with a separate classification branch. Our experiments to learn tracking and identification from the same backbone with an additional head were less successful, achieving an accuracy of just 41%. This difference in performance could be explained by small training data set and the fact that the tracking model with the additional classification head was required to learn two competing tasks simultaneously. When focusing on tracking, the model may discard unnecessary information that might be helpful for individual identification. Similarly, tracking performance decreased when giving more weight to the identification head. This problem has been observed in the past when combining object detection with human-object interaction detection (Chen et al., 2021). The combined approach might work in applications with more annotated data, and the benefit of having only one backbone and different lightweight heads for downstream tasks could be achievable in this context. The classification branch can be used in cases where all individual identities are known. If there are unknown individuals in the dataset, open-set classification approaches (e.g. via Deep metric learning) (Vidal et al., 2021) are a more promising pathway as they do not require a predefined set of individuals.

### Future directions

The flexibility of the model allows the extension of PriMAT to incorporate other tasks such as instance segmentation (Wang et al., 2020) and ultimately interaction classification (Liu et al., 2021) between objects when those interactions are detectable in single frames. The classification branch can be used for individual identification but also to classify actions performed by the individuals. In the computer vision community, many models developed for spatio-temporal action recognition, interaction recognition or gaze following build on bounding boxes as intermediate representations (Chong et al., 2020; Woo et al., 2022; Feng et al., 2023; Ryan et al., 2024).The very recent PanAf500 dataset (Brookes et al., 2024) contains nine activity classes, such as sitting, walking, climbing up or down and would be a suitable benchmark. While our primary focus was on two species of nonhuman primates (Assamese macaques and redfronted lemurs), we observed similar applicability in videos of other nonhuman primate species (including Barbary macaques, Guinea baboons, chimpanzees and gorillas) and expect the model to generalize well to other animal species with little additional annotation effort.

## Supporting information

Images of qualitative results; Results of hyperparameter search

## Acknowledgements

This project was funded by the Deutsche Forschungsgemeinschaft (DFG, German Research Foundation) - Project-ID 454648639 - SFB 1528. We acknowledge funding by the Leibniz Association through an Audacity Grant from the Leibniz ScienceCampus Primate Cognition (W45/2019 – Strategische Vernetzung) and by the Deutsche Forschungsgemeinschaft (DFG, German Research Foundation), Grant/Award Number: 254142454 / GRK 2070. We thank the Department of National Parks, Wildlife, and Plant Conservation (DNP) and the National Research Council of Thailand (NRCT) for research permission. This work used the Scientific Compute Cluster at GWDG, the joint data center of Max Planck Society for the Advancement of Science (MPG) and University of Göttingen.

## Footnotes

1 https://github.com/ecker-lab/PriMAT-tracking

2 https://github.com/microsoft/VoTT

3 https://www.cvat.ai

4 https://github.com/microsoft/VoTT

