## Supplementary material for "PriMAT: A robust multi-animal tracking model for primates in the wild": Images of qualitative results; Results of hyperparameter search

### Qualitative comparison

The quantitative results from the comparison between keypoint tracking and bounding box tracking can also be seen qualitatively. We see that the keypoint skeletons of the lemurs are almost all incomplete, some of them are distributed across two lemurs and sometimes two skeletons are found within one individual. Some lemurs remain undetected by the keypoint model (Figure 1).

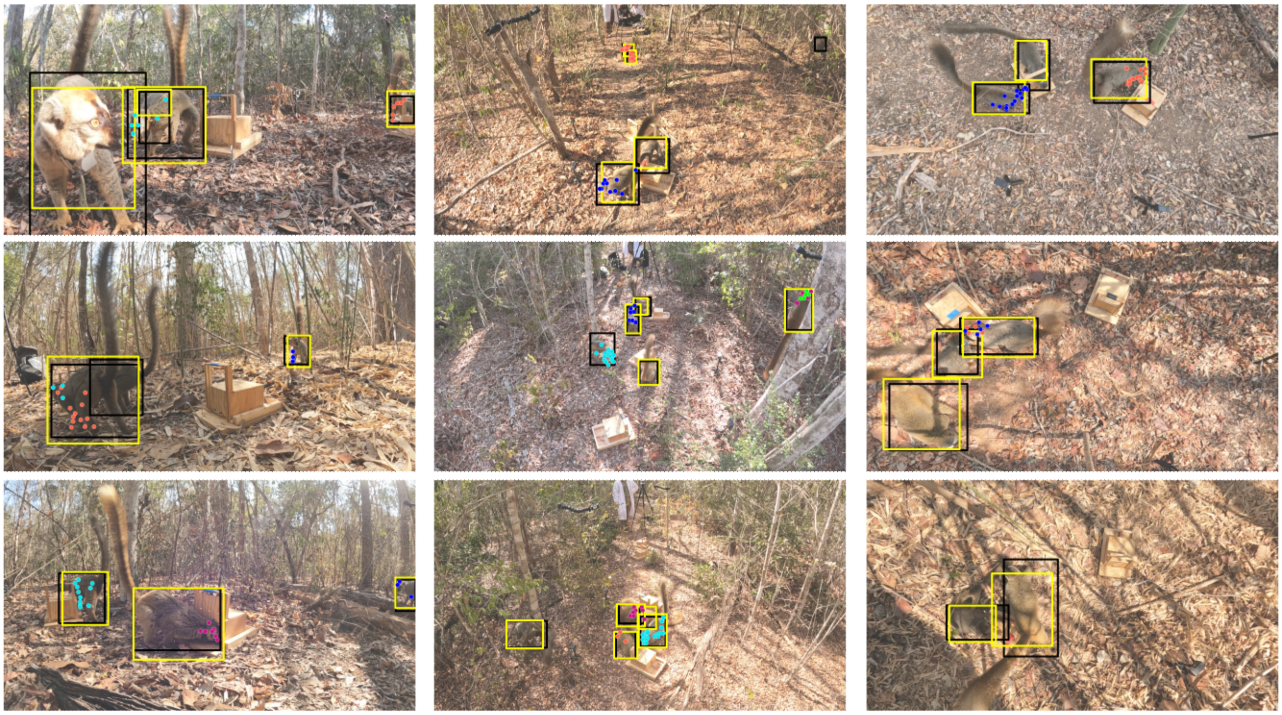

Figure 1: Qualitative comparison between PriMAT and DLC. Ground truth bounding boxes are black, PriMAT bounding boxes are yellow. The different colors of the keypoints refer to different detected individuals.

### Transfer to other nonhuman primate species and settings

Qualitative examples show that our model is able to detect most nonhuman primates in those videos without further training (Figure 2A). However, there are some clear limitations. The low-resolution videos of PanAf500 are very different to the recordings of lemurs and macaques and some individuals are not detected. When individuals get close to the camera, sometimes several detections within the same individual are made. And lastly, some objects that were not present in the training data are sometimes detected as primates (Figure 2B). We selected and annotated 100 frames for each of the three newly introduced settings. We excluded videos from which we had taken our qualitative examples, such that the models would not overfit to the specific video. Starting from our model trained on lemurs and macaques, we finetuned each model for 50 epochs with the respective annotated frames. Afterwards, the models were able to avoid the previously described error cases (Figure 2C).

### Tracking

**Backbone and heads.** The backbone processes the input images $I \in [0, 255]^{W_0 \times H_0 \times 3}$ into feature maps $F \in \mathbb{R}^{W \times H \times C}$, where $W = W_0/4$, $H = H_0/4$ (in our case $W \times H = 272 \times 152$ and $C = 512$). We use HRNet (Sun et al., 2019) as a backbone. Four independent heads process the resulting feature map, each with the same width and height, but with a different number of channels and a different loss, depending on their task. The four heads are:

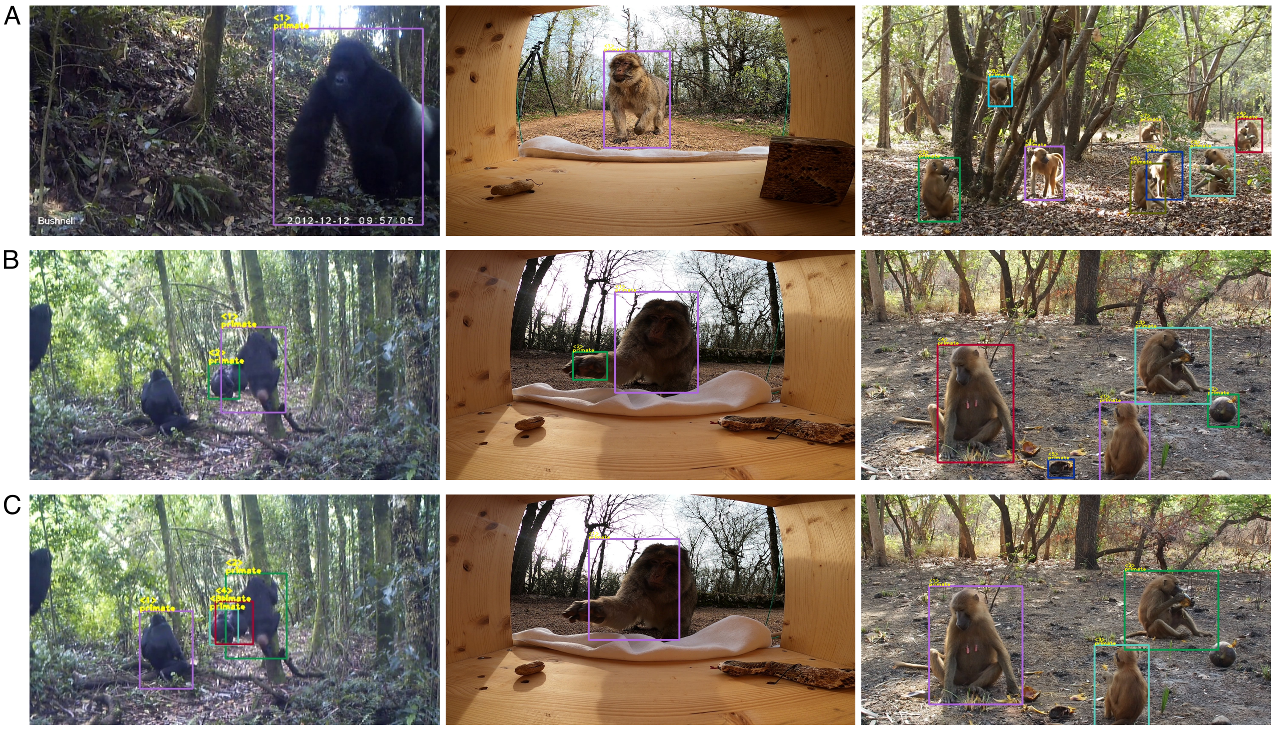


Figure 2: Qualitative examples of our model trained jointly on macaques and lemurs, and applied without further training to different species and settings (from left to right: PanAf500 dataset with gorillas and chimpanzees (Brookes et al., 2024), recordings of a decision-making test with Barbary macaques in Rocamadour, France (Bosshard et al., 2024), Guinea baboons ranging near the CRP Simenti Field Station in the Niokolo Koba National Park in Senegal (Fischer et al., 2017)). (A) positive examples without further training on the target dataset. (B) Error cases: missing individuals, additional detection for limbs when approaching the camera, detection of other objects as primates. (C) After retraining with 100 annotated samples from the specific dataset (selected from different videos), the error cases can be circumvented.

- Center heatmap (output dimensionality $W \times H \times cl$): This head predicts for each pixel how likely it is to be the center of an object. We adapted the architecture to be a multi-class model by having one center heatmap per class. $cl$ is given by the number of different object classes that should be tracked; in the lemur case $cl = 2$, for tracking lemurs and feeding boxes and $cl = 1$ for the macaque model, as only macaques are being tracked.
- Bounding box size ($W \times H \times 2$): This head predicts for each pixel the width and height of a hypothetical bounding box if there is a center point at this pixel. The two channels correspond to the width and height of the respective bounding box.
- Bounding box offset ($W \times H \times 2$): The feature map $F$ has a resolution four times smaller than the input image size. Therefore, each pixel on the feature map corresponds to $4 \times 4$ pixels in the original image. This head predicts a small offset for each predicted bounding box to correct for the effect of down-sampling.
- Re-identification (ReID) features ($W \times H \times 128$): The ReID head generates a numeric representation as a vector for each detected object. This vector encodes the individual's appearance and is later used as additional information for the association of individuals over time.

For more information about the heads and loss functions, please refer to the detailed description in FairMOT (Zhang et al., 2021).

**Association of detections.** The general idea for the association is the same as in FairMOT (Zhang et al., 2021): Initially, the model identifies objects from the local maxima in the center heatmap of the first frame, creating individual tracks. In subsequent frames, it generates new detections and ReID features, comparing them to existing tracks for similarity. When they match, the detections are added to the corresponding tracks, while unmatched detections are considered potential candidates for new tracks.

The similarity between existing tracks and new detections is assessed using two criteria, appearance and location. Appearance similarity is calculated via cosine similarity of the ReID features, whereas intersection-over-union (IoU) is the measure for similarity in location. Additionally, motion information is modeled using a Kalman filter which predicts the expected location of each bounding box based on velocity in the past frames. The result is a distance matrix between the set of existing tracks (i.e. past detections) and the set of detections in the current frame.

While FairMOT followed a more complicated procedure that involved a sequential application of the two distance measures, we simplified the process as earlier experiments showed that it performed at least equally well. In our model, the final distance matrix is calculated as a linear combination between the IoU distance matrix and the distance matrix obtained from the appearance features. We found a weight of 0.8 for the IoU distance and 0.2 for the appearance feature distance to work well in our applications. The optimal assignment of new detections to existing tracks is obtained using Hungarian matching (Kuhn, 1955). Unmatched tracks are retained for 90 frames to facilitate re-detection after occlusion periods (see Table 1).

**Inter-annotator reliability** For quality control of the annotations used for evaluation, we assessed the consistency between the labels produced by two human annotators on a subset of three videos of the macaque dataset. We matched the annotated bounding boxes using Hungarian matching and calculated the average intersection over union across all boxes. The average IoU score per video was over 0.8 for the three videos.

**Hyperparameters.** The motion patterns and appearance of monkeys and lemurs differ from those of humans. Therefore, PriMAT contains elements that are specifically tailored to videos of primates in the wild. We described these in Section **??** and evaluated how they help to improve performance. We tested hyperparameters to show their influence on model performance (Table 1).

- Association method: Comparing regular association via Kalman filter and our adaptation for fast motion and jumps, which additionally compares each new detection to the last visible position of the track, if regular assocation does not find a match.
- Confidence threshold: How confident should the model (i.e. the center heatmap) be to propose a detection at a certain location?
- Detection threshold: How confident should the model be with a detection to start a new track if it was not matched to any existing track?
- Association threshold: We combined IoU and cosine similarity by a linear combination of both matrices. The Hungarian matching returns the association with the highest similarity values between existing tracks and detections. If the best match lies below this association threshold, we leave them unmatched.
- When associating new detections with existing tracks, we calculate a similarity measure from IoU (location) and cosine similarity of the ReID features (appearance). The proportion IoU parameter determines how much importance is giving to the location. The importance to appearance is automatically one minus the value for IoU.

The optimal values differ slightly from the ones reported in Table **??** as we did the extensive hyperparameter search only on the best performing pretraining dataset for each species (ImageNet for lemurs, MacaqueCopyPaste for macaques).

**Robustness with smaller samples sizes.** We ran experiments on how the size of the dataset influences performance. For this, we split our datasets containing 500 images into disjoint $5 \times 100$ and the same $2 \times 200$ frames mentioned above and trained PriMAT models with them. For the macaques, the five models had a mean of 64.3 HOTA and a standard deviation of 1.28 (ranging from 62.5 to 65.6).

Table 1: Comparison of tracking performance on the 12 validation video sequences for each, lemurs and macaques. a) An additional association step to prevent identity switches after jumps or fast motion improved performance for lemurs, but not for macaques, where the validation videos did not contain any jumps. b) The confidence threshold for pedestrian tracking is recommended at 0.4. A reduced detection threshold showed improved results for both lemurs and macaques. c) The detection threshold needs to be surpassed for an unmatched detection to start a new track. d) The association threshold is the minimum amount of similarity required between a detection and a track to form a match. e) The proportion IoU parameter manages the relative importance between location and appearance for matching detections and tracks. f) The track buffer determines for how long unmatched tracks are being stored to find a matching detection e.g. after a period of occlusion. Our videos had a frame rate of 30 and any track buffer over 10 frames performed equally well.

| | **Lemurs** | | | **Macaques** | | |
|---|---|---|---|---|---|---|
| | **HOTA** | **MOTA** | **IDF1** | **HOTA** | **MOTA** | **IDF1** |
| **a) Association** | | | | | | |
| Regular | 68.2 | 78.1 | 84.9 | **66.6** | **75.5** | **83.7** |
| Fast motion | **70.3** | **81.5** | **88.1** | 64.5 | 74.0 | 80.6 |
| **b) Confidence threshold** | | | | | | |
| 0.01 | **70.3** | 81.5 | **88.1** | 64.1 | 70.7 | 78.3 |
| 0.02 | 69.8 | **81.8** | 87.2 | 65.8 | 73.8 | 82.0 |
| 0.04 | 69.4 | 81.2 | 86.6 | **66.6** | 75.5 | **83.7** |
| 0.1 | 69.0 | 81.2 | 86.7 | 65.7 | **76.7** | 82.2 |
| 0.2 | 68.8 | 80.0 | 86.7 | 65.6 | 75.8 | 83.0 |
| 0.4 | 66.9 | 78.6 | 84.4 | 63.0 | 72.9 | 78.8 |
| **c) Detection threshold** | | | | | | |
| 0.4 | **70.3** | 81.4 | 88.0 | 66.1 | 74.5 | 82.7 |
| 0.5 | **70.3** | **81.5** | **88.1** | **66.6** | 75.5 | **83.7** |
| 0.6 | 70.2 | 81.2 | 87.9 | 66.5 | **76.3** | 83.6 |
| **d) Association threshold** | | | | | | |
| 0.7 | **70.3** | **81.5** | **88.1** | **66.6** | **75.5** | **83.7** |
| 0.8 | 69.5 | 80.1 | 86.8 | 65.4 | 74.0 | 82.0 |
| 0.9 | 68.8 | 78.1 | 86.4 | 63.5 | 72.9 | 78.9 |
| **e) Proportion IoU** | | | | | | |
| 0 | 68.3 | 77.8 | 85.3 | 61.5 | 72.7 | 75.6 |
| 0.1 | 69.1 | 79.3 | 86.8 | 63.7 | 73.5 | 79.0 |
| 0.2 | 69.3 | 80.0 | 86.8 | 65.0 | 74.6 | 81.1 |
| 0.3 | 69.6 | 81.1 | 87.1 | 64.9 | 75.5 | 80.5 |
| 0.4 | 69.6 | 81.0 | 87.3 | **66.6** | 75.5 | **83.7** |
| 0.5 | 70.2 | 81.4 | 87.8 | 65.3 | 75.9 | 81.0 |
| 0.6 | **70.3** | **81.5** | **88.1** | 65.9 | 76.6 | 82.5 |
| 0.7 | 70.2 | 81.4 | 88.0 | 65.9 | 77.4 | 82.6 |
| 0.8 | 70.0 | 81.0 | 87.8 | 65.2 | **77.5** | 80.7 |
| 0.9 | 69.9 | 80.8 | 87.6 | 65.1 | 77.4 | 80.6 |
| 1 | 69.9 | 80.7 | 87.7 | 64.9 | **77.5** | 80.2 |
| **f) Track buffer** | | | | | | |
| 0 | 68.6 | 78.9 | 85.7 | 63.3 | 76.5 | 75.6 |
| 1 | 68.6 | 78.9 | 85.7 | 63.3 | 76.5 | 75.6 |
| 5 | 69.5 | 79.8 | 87.0 | 65.4 | **76.9** | 80.2 |
| 10 | 70.2 | 81.2 | 87.9 | 66.5 | 76.8 | 83.6 |
| 30 | **70.3** | **81.5** | **88.1** | 66.5 | 76.1 | 83.5 |
| 90 | **70.3** | **81.5** | **88.1** | **66.6** | 75.5 | **83.7** |
| 180 | **70.3** | **81.5** | **88.1** | **66.6** | 75.5 | **83.7** |

The two models trained with 200 samples had HOTA values of 62.5 and 64.8. The model trained with all 500 samples had a HOTA of 66.1. For the lemurs, the five models had a mean of 62.4 HOTA and a standard deviation of 1.08 (ranging from 61.0 to 63.6). The two models trained with 200 samples had

Table 2: Showing HOTA scores for lemurs and macaques when the model is trained with a subset of the training data. Subscripts refer to disjoint subsets.

| Sample size$_{split}$ | HOTA (lemurs) | HOTA (macaques) |
|---|---|---|
| 500 | 70.3 | 66.1 |
| $200_1$ | 65.7 | 64.8 |
| $200_2$ | 66.5 | 62.5 |
| $100_1$ | 63.3 | 65.6 |
| $100_2$ | 61.0 | 64.1 |
| $100_3$ | 62.2 | 65.5 |
| $100_4$ | 61.8 | 63.9 |
| $100_5$ | 63.6 | 62.5 |

HOTA values of 65.7 and 66.5. The model trained with all 500 samples had a HOTA of 70.3 (Table 2). This shows that the training of the bounding box models converges to stable solutions which do not show a large variance between different datasets.

### Individual identification

**Majority voting.** We get framewise predictions for each individual. On many of the frames the individual is difficult to identify, however, when the collar or the face is visible, the model is more confident in its predictions, and can assign a high probability to one individual. To take a final prediction for the whole track, we decided to take a majority vote across all predictions and tested different strategies. (1) **Maximum**: Finding the moment with the highest probability voting for one individual and taking that individual. (2) **Average**: Averaging the probability of all individuals across the track and taking the individual with the highest average probability. (3) **Count**: Counting how often an individual was predicted with a minimum confidence threshold and taking the individual with the highest count. (4) **Two thresholds**: Working with two thresholds: Counting every individual that was predicted over the lower threshold, and multiplying with 100 every appearance over the higher threshold. (5) **Weighting**: Exponentially weighting the predictions. The function $w = \exp(9.2 \times (p - 0.5))$ where $p$ is the probability assigned to the highest individual has the property that it assigns a weight of 100 if the model assigns the probability of 1 to an individual, and a weight of 1, when the probability is 0.5. It exponentially adds more weights to higher confidence predictions. Counting with thresholds and exponential weighting outperform the other methods (see Table 3).

Table 3: Performance of different majority voting strategies on individual identification.

| Strategy | Threshold(s) / Weights | Accuracy |
|---|---|---|
| Maximum | | 0.684 |
| Average | | 0.771 |
| Count | 0.2 | 0.793 |
| Count | 0.5 | 0.826 |
| Count | 0.7 | 0.772 |
| Two thresholds | 0.5 / 0.99 | 0.826 |
| Weighting | $\exp(9.2 \times (p - 0.5))$ | 0.837 |
